# The evolutionary trajectory of drosophilid walking

**DOI:** 10.1101/2021.09.29.462444

**Authors:** Ryan A. York, Luke Brezovec, Jenn Coughlan, Steven Herbst, Avery Krieger, Su-Yee Lee, Brandon Pratt, Ashley Smart, Eugene Song, Anton Suvorov, Daniel R. Matute, John C. Tuthill, Thomas R. Clandinin

## Abstract

Neural circuits must both execute the behavioral repertoire of individuals and account for behavioral variation across species. Understanding how this variation emerges over evolutionary time requires large-scale phylogenetic comparisons of behavioral repertoires. Here, we describe the evolution of walking in fruit flies by capturing high-resolution, unconstrained movement from 13 species and 15 strains of drosophilids. We find that walking can be captured in a universal behavior space, the structure of which is evolutionarily conserved. However, the occurrence of, and transitions between, specific movements have evolved rapidly, resulting in repeated convergent evolution in the temporal structure of locomotion. Moreover, a meta-analysis demonstrates that many behaviors evolve more rapidly than other traits. Thus, the architecture and physiology of locomotor circuits can both execute precise individual movements in one species and simultaneously support rapid evolutionary changes in the temporal ordering of these modular elements across clades.

## Introduction

A central goal of neuroscience is to understand how circuits shape behavior, efforts that have greatly advanced our understanding of neural mechanisms using a relatively small number of model systems. However, how the structure and function of neural circuits can support the diversification behavioral repertoires over evolutionary time is incompletely understood. To understand how circuits can evolve requires establishing model clades, groups of related species displaying both behavioral diversity and having the requisite tools to dissect circuit and computational mechanisms (Katz 2019; Gallant & O’Connell 2020; Jourjine & Hoekstra 2021). New methods in behavioral measurement (Pereira et al. 2020, Mathis et al. 2020) and statistical analysis (von Ziegler et al. 2021, York et al. 2021) have made it possible to collect and study many aspects of behavior at large scale in many clades. Similarly, broadly sampled and time-resolved molecular phylogenies have become increasingly available for many groups of animals (Piel et al. 2009). Here, we demonstrate that the combination of such high-resolution, quantitative behavioral analyses with robust phylogenetics can reveal rich patterns of behavioral evolution comparable to those seen for other traits.

We focus on the evolution of walking, a critical element of many behaviors, in fruit flies of the genus *Drosophila.* Fruit flies are nearly unique in being both a genetic *and* an evolutionary model system, representing a broad range of life histories and ecological contexts that provide strong bases for inter- and intra-specific comparisons (Markow & O’Grady 2005; Hales et al. 2015; Markow 2015; Jezovit et al. 2017). Moreover, the structure of fruit fly walking has been well resolved in the model species *D. melanogaster* (Berman et al. 2014, Katsov et al. 2017, Tao et al. 2019) and recent work has characterized the functions of a number of critical neuron types that determine the initiation of walking, turning and stopping (Bidaye et al. 2014; Carreira-Rosario et al. 2018; Namiki et al. 2018; Cande et al. 2018; Feng et al. 2020). Here we develop an approach to quantitatively compare the structure of walking across fly species and strains. We find that this well-constrained example of motor control can evolve surprisingly rapidly, with closely related strains diverging, and distantly related species converging on similar temporal patterns of locomotor movements. These results demonstrate that behavioral variation can emerge from changing the temporal sequence of individual, modular movements, thereby identifying potential neural mechanisms of locomotor evolution.

## Results

### High-throughput measurement of drosophilid walking

To analyze the structure of naturalistic walking in fruit flies we developed the Coliseum, a novel apparatus for behavioral experiments in *Drosophila* (**Figure 1A**), and flyvr software (Methods) for the acquisition and analysis of data from the Coliseum (**Figure 1B**). The Coliseum uses a fly-initiated dispenser to introduce flies onto a darkened ~1m x 1m platform under infrared illumination (**Figure 1A**; Methods). As flies freely explore the apparatus, a camera attached to a stepper-motor system tracks and updates the camera position with animal movement (100 Hz), yielding high-quality video and positional data in realtime (**Figure 1B**). We used the Coliseum to measure the trajectories of 1,030 individuals from 13 globally distributed *Drosophila* species with diverse distributions from cosmopolitan (e.g. *D. melanogaster, D. simulans),* temperate (e.g. *D. virilis, D. persimilis),* tropical (e.g. *D. yakuba, D. mauritiana),* and desert (e.g. *D. arizonae)* habitats (Jezovit et al. 2017). In addition, to detect traits that can undergo very rapid evolutionary changes, we analyzed 15 wild-derived strains of *D. melanogaster* from its ancestral distribution in Africa (**Figure 1C**).

**Figure 1.**
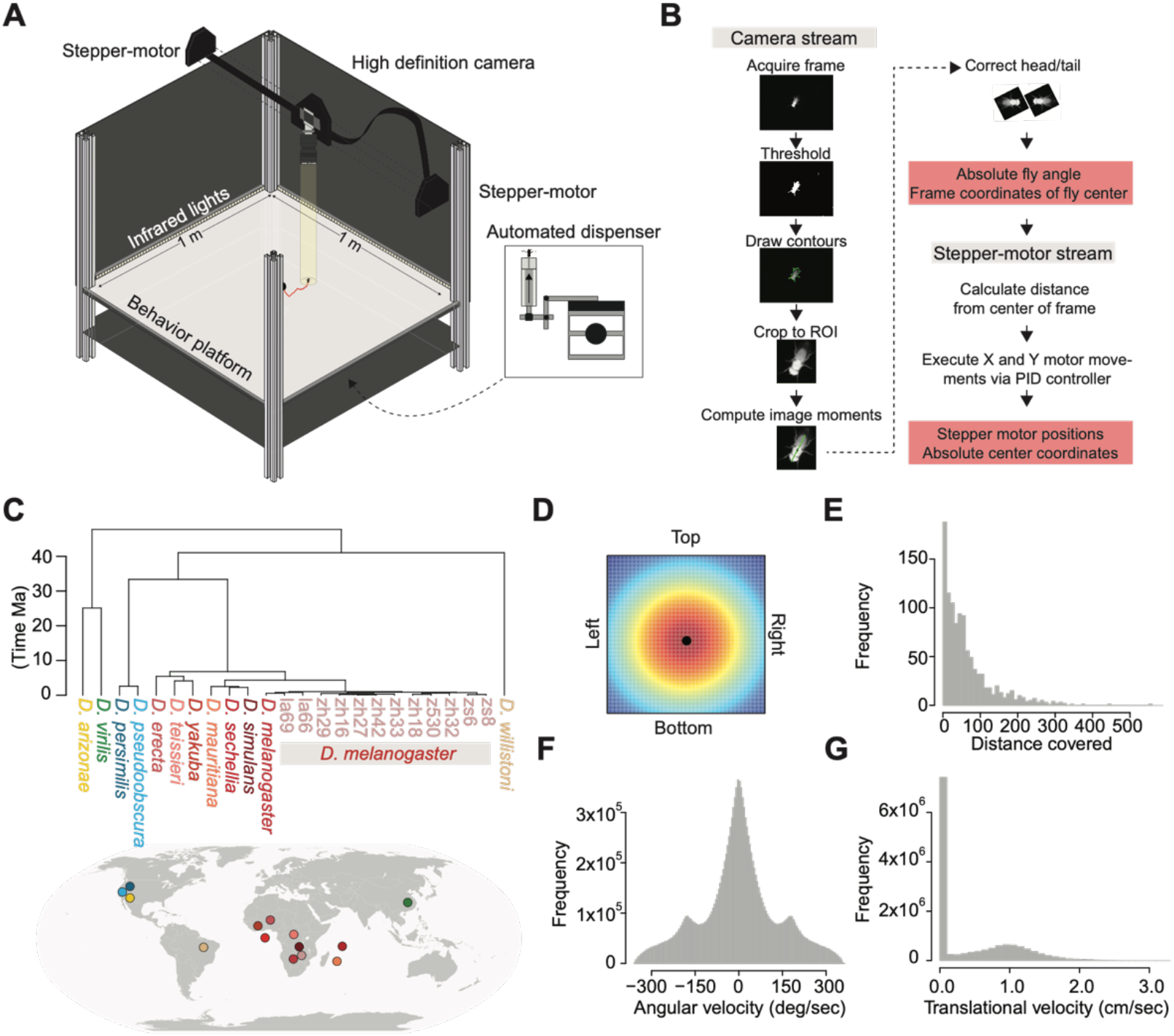
High-throughput measurement of locomotion across *Drosophila*. **(A)** Schematic of the Coliseum. Flies are introduced onto an IR illuminated behavior platform by an automated dispenser. A high-definition camera is suspended above on a stepper-motor system that updates the camera’s position as the fly moves. The visual field of the camera is indicated by the yellow cylinder tracking the fly’s trajectory (marked by the red path). **(B)** Workflow for acquiring fruit fly position in the Coliseum, extracting orientation, and updating camera position via stepper motors. **(C)** (above) Time-calibrated phylogeny of the species analyzed. (below) The approximate global locations of the ancestral populations for each species or strain represented in the phylogeny (coded by color). **(D)** Probability density function of fly position in the Coliseum, computed using the positions of all trials in the data set. The dispenser hole location is indicated by the black dot. **(E-G)**, The distributions of distance covered (**E**), angular velocity (**F**), and translational velocity (**G**) encompassed by the full locomotor data set.

Comparing their relationships using a fossil-calibrated phylogeny established that these species and strains represent ~40 million years of evolutionary history (**Figure 1C**). A variety of species and clade-level relationships were examined, including recent evolutionary diversifications in the *D. melanogaster* clade (crown age = 41 MYA; minimum branch length = 0.12 MYA; **Figure 1C**), facilitating a phylogenetically thorough exploration of the evolution of walking. Overall, flies uniformly explored the Coliseum (**Figure 1D**) and displayed a variety of trajectory lengths and durations (**Figure 1E**) within the typical kinematic range of adult *Drosophila* (**Figure 1F-G**) (Katsov et al. 2017, Tao et al. 2019, DeAngelis et al. 2019). These measurements varied as a function of species and strain (**Figure S1A-D**), highlighting diversity in these first order statistics across genotypes. The Coliseum is therefore a powerful tool for measuring naturalistic walking and can be used to obtain high-resolution data from diverse strains and species.

### A common space captures drosophilid walking

The first order statistics of walking display substantial temporal structure on the timescale of hundreds of milliseconds (Berman et al. 2014, Katsov et al. 2017, Tao et al. 2019, DeAngelis et al. 2019). To capture this structure, we employed the TREBLE framework (**Figure 2A**; York et al. 2021) to create a common behavior space encompassing all individual flies from all strains and species (**Figure S1J**). Briefly, kinematic parameters (**Figure 2A ii**) were measured from the raw fly trajectories (**Figure 2A iv**), binned into windows that reflected the temporal structure of walking (see Methods; **Figure 2A iii**, **S1E-I**), and embedded into a 2-d behavior space via non-linear dimensionality reduction (McInnes et al. 2018; **Figure 2A iv**). The resulting space separated kinematic parameters along identifiable axes (**Figure 2B**) corresponding to recognizable behaviors (**Figure 2C**, **S1L-N**). To identify stereotyped movement through the space, we calculated the mean path of all flies through every point in behavior space, producing a vector map in which large vectors correspond to similar movements performed by many animals. This analysis revealed common pathways linking specific behaviors that were associated with high covariation (**Figure 2D-E**). TREBLE therefore produced a stereotyped, continuous behavior space encompassing the breadth of drosophilid walking kinematics.

**Figure 2:**
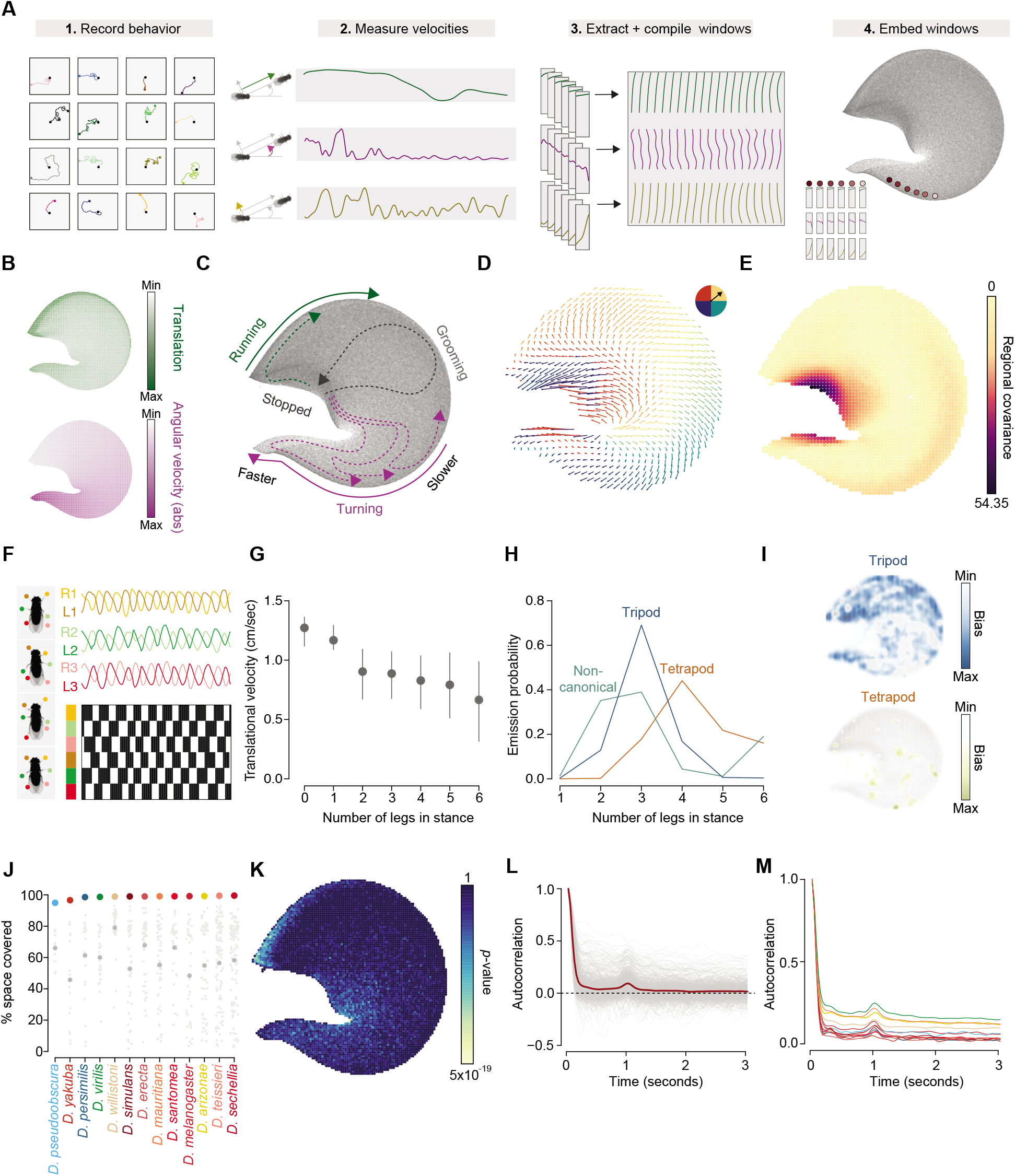
Defining a universal walking behavior space. **(A)** The workflow for the TREBLE framework. **(B)** The distributions of translational (top) and angular (bottom) velocities as a function of position in behavior space. **(C)** The structure of behavior space annotated with behavioral states and pathways in between. **(D)** The mean vector field of behavior space. Arrow angle and length indicate the direction and magnitude, respectively, of probabilistic movement between points. The angle degree is also denoted by color, corresponding the circle plotted above. **(E)** The distribution of temporal covariance as a function of position in behavior space. Darker colors correspond to regions in which points have similar covariance values (i.e. possess more stereotyped patterns of movement through behavior space). **(F)** The measurement of gait parameters from videos of freely walking flies. Positions of the fly’s 6 tarsi are acquired for each frame (fly images on left) and are then egocentrically aligned and converted to phase (panel top right) from which swing-stance estimates are made (panel bottom right). Sample distributions are from *D. melanogaster.* **G)** The distribution of translational velocity as a function of number of legs in stance across all genotypes. **(H)** Emission probabilities corresponding to number of legs in stance as a function of HMM state. **(I)** The distributions of tripod- and tetrapod-biased densities in behavior space. Darker color corresponds to more bias toward the given gait type. **(J)** The percent of behavior space covered by pure species in the dataset. Individual trial percentages are denoted per-species as grey points, the mean of which is represented by the larger dark grey point. **(K)** The distribution of per-bin significance in variance of occurrence across species (measured by Kruskal-Wallis test *P-*values calculated across all species, Bonferroni corrected). Lighter colors correspond to increasingly significant *P*-values, representing substantial variation in occurrence between species. **(L-M)**, Autocorrelation functions of behavior space position over a 3 second span for all individuals (**L**; mean function denoted by red line) and species means (**M**).

Distinct gait patterns are associated with variation in kinematics over time in *D. melanogaster* (Mendes et al. 2013, Wosnitza et al. 2013, Isakov et al. 2016, Pereira et al. 2019, DeAngelis et al. 2019, Chun et al. 2021). To explore whether other species displayed similar patterns, we identified the positions of all six limbs for each species and strain and extracted the phase and swing-stance state at each frame (**Figure 2F**; Methods). As expected from *D. melanogaster* (Mendes et al. 2013, Wosnitza et al. 2013, Isakov et al. 2016, Pereira et al. 2019, DeAngelis et al. 2019, Chun et al. 2021), we found that the number of legs in stance was speed dependent across all genotypes wherein velocity decreased as the number of legs in stance increased (**Figure 2G**). We next fit Hidden Markov Models (HMMs) predicting the number of legs in stance given a varying number of hidden states (Methods; **Table S1**). Notably, a model with three hidden states representing tripod, tetrapod, and non-canonical gaits fit the data the best, again matching *D. melanogaster* (Isakov et al. 2016, Pereira et al. 2019, DeAngelis et al. 2019) (**Figure 2H**). While each state displayed a characteristic distribution across behavior space (**Figure 2I**), tripod gait was associated with a much broader distribution in comparison to the other states, corroborating findings that tripod gait can be employed broadly across walking speeds (Chun et al. 2021). These observations suggest that the relationship between gait and kinematics across *Drosophila* is similar to that seen in *D. melanogaster.* Moreover, position in behavior space maps onto identifiable gait patterns that are conserved across species.

While the above observations suggest conservation of the overall structure of behavior space, it is possible that locomotor evolution led to the development of unique kinematic combinations in subsets of lineages. Such changes would result in species- or clade-specific clustering within behavior space. To test this possibility, we analyzed the amount of behavior space covered by each species and observed that each of them explored at least 95% of the space (**Figure 2J**). However, there was significant variation in the amount of space covered by individual flies from each species (Kruskal-Wallis test; *P* = 4.2 × 10^-6^; **Figure 2J**), reflecting variation in the amount of time spent in specific regions (**Figure 2K**, **3A**) and mirroring the observed differences in first-order kinematic parameters (**Figure S1A-D**). Furthermore, variation in these traits was greater between groups than within, indicating that these differences arise from distinct behavioral differences between genotypes rather than context (**Figure S2A-B**; Methods).

**Figure 3:**
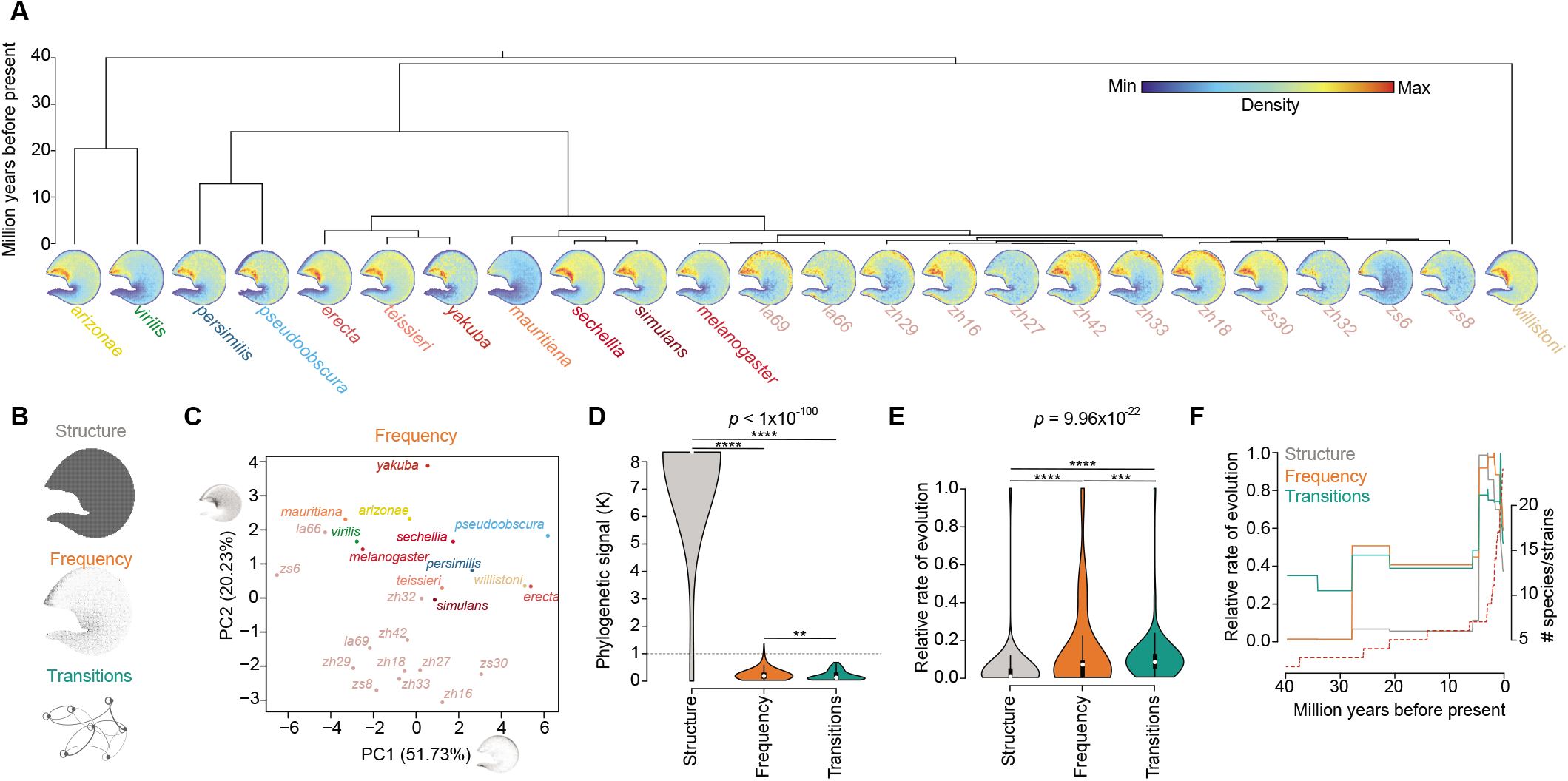
The dynamics of locomotor evolution across *Drosophila*. **(A)** The phylogenetic distribution of behavior space density maps across all species/strains. **(B)** Cartoons of the three traits considered in the comparative phylogenetic analyses: *structure, frequency,* and *transitions.* **(C)** *Frequency* morphospace. Variance explained by the first two PCs are denoted in the axes. PC loadings are represented by the grayscale density maps of behavior space on the x and y axes. **(D)** Violin plot of the distribution of phylogenetic signal across behavior space for *structure, frequency,* and *transitions* (see Methods; Kruskal-Wallis test followed by post-hoc Dunn’s test) \*\*\*\**P*<0.0001, \*\*\**P*<0.001. **(E)** Violin plot of the comparison of relative rate of evolution measurements across all nodes in the phylogeny for all three traits (Kruskal-Wallis test followed by post-hoc Dunn’s test) \*\*\*\**P*<0.0001, \*\**P*<0.001. **(F)** Mean relative rates of evolution over time for each trait (computed in 0.1 million year windows). Species accumulation is indicated by the dotted red line.

Are these differences in frequency mirrored by changes in the sequencing of movements? Since position in behavior space represents an animal’s current movement at a given time point, we reasoned that the autocorrelation of position could be used as a proxy for measuring the temporal structure of behavior. We found a common pattern across all flies in which movements were tightly correlated over several hundred milliseconds, declined rapidly, and rebounded briefly around a second (**Figure 2L**). However, while each individual species showed this same autocorrelation pattern, they also differed in their baseline levels of autocorrelation (**Figure 2M**). These lines of evidence suggest that the frequency and sequencing of individual movements, but not the range of kinematic parameter combinations, have seen increased differentiation over evolutionary time.

### Tempo and dynamics of the evolution of walking

To explore the evolutionary dynamics of these patterns, we leveraged a fossil-calibrated genome-scale phylogeny of *Drosophila* (Suvorov et al. 2021; Methods) to perform a series of comparative phylogenetic analyses (**Figure 3A**). In phylogenetic studies of traits such as morphology, the evolutionary diversification of multiple components of a phenotype can be mapped, providing insights into patterns of developmental constraint, trait hierarchy, and phenotypic correlations (Adams & Collyer 2009; Felice et al. 2018). In these frameworks, multi-dimensional traits are often represented in a lower dimensional morphological space (‘morphospace’) that represents the range of trait values possible across the species analyzed. Given that the TREBLE behavior space captures the full range of a behavioral trait (walking kinematics), we wondered whether the structure and dynamics of this space might be studied in a fashion analogous to morphological traits.

We considered three behavioral traits: *structure* (% of behavior space explored), *frequency* (probability of occurring in specific regions), and *transitions* (probability of sequencing between specific regions; Methods) (**Figure 3B**). This hierarchical description of behavior represents, in order, the movements that are possible, how often they are performed, and how often certain movements precede others. We created a ‘behavioral morphospace’ for these three traits using principal component analysis (PCA) (**Figure 3C**, **S3A-C**; Methods). The first two components of the morphospace for *structure* separated species by the amount of behavior space covered (**Figure 2J**, **S3A**). On the other hand, *frequency* and *transitions* morphospaces yielded more unexpected results. The first two components of the *frequency* morphospace together explained 72% of variation in the trait (**Figure 3C**, **S3B**). Comparing the distribution in morphospace to the phylogenetic patterns of space occupancy (**Figure 3A**) demonstrated that PC1 separated high and low activity states (**Figure 3C**; 51.73% variance explained). Conversely, PC2 separated *D. melanogaster* strains and *D. melanogaster*-like species from all others (**Figure 3C**; 20.23% variance explained). The first two components of the *transitions* morphospace explained 81% of variation (**Figure S3C**). In this case, however, PC1 separated *D. melanogaster* strains from the rest of the species (66.97% variance explained) while PC2 represented behavioral differences associated with fast movements (**Figure S3C**; 13.75% variance explained). These observations suggest that, in the cases of *frequency* and *transitions,* there is a substantial degree of trait variation not associated with phylogenetic relationships providing initial evidence of behavioral convergence across distantly related species.

We then estimated the evolutionary lability of these traits by calculating the distribution of phylogenetic signal for each (Methods). Phylogenetic signal varied as a function of the behavioral hierarchy (**Figure 3D**). *Structure* was extremely conserved (mean Blomberg’s *K* = 8.32, sd = 2.14) while the mean values of both *frequency* (mean Blomberg’s *K* = 0.24, sd = 0.14) and *transitions* (mean Blomberg’s *K* = 0.16, sd = 0.11) were well below 1, suggesting rapid evolution of these traits independent of phylogenetic relationships. Furthermore, the phylogenetic signal distributions across all three traits varied significantly (Kruskal-Wallis test: *P* < 1×10^-100^) with *transitions* having a distribution significantly lower than *frequency* (Dunn’s test: *P* = 0.0088; *z* = 2.37), suggesting a particularly rapid tempo of evolution in the sequencing of behavior across the *Drosophila* tree.

To assess the temporal dynamics of these patterns, we estimated ancestral states with variable-rate models of evolution and examined variation in morphospace occupation over time. Across the full phylogeny, *transitions* evolved at the fastest rate (mean normalized relative evolutionary rate = 0.14; Methods) followed by *frequency* (rate = 0.11) and *structure* (rate = 0.05). Each trait had a distinct distribution of rates across the phylogeny (**Figure 3E**) that varied significantly from the others and corroborated the phylogenetic signal analyses. Accordingly, the rate of evolution of *structure* largely mirrored the accumulation of species (**Figure 3F**, **S3D**), increasing in rate in parallel with the diversification of the *D. melanogaster* clade over ~6.7 Ma. On the other hand, *frequency* and *transitions* displayed multiple epochs of increases and decreases in rate across the phylogeny (**Figure 3F**, **S3D**).

To study the rate of behavioral trait diversification, we examined how each morphospace was populated over evolutionary time (**Figure S3E-G**; Methods). We found that the full range of *structure* morphospace was sparsely sampled (**Figure S3F**), covering just 19% of phenotypic values and occurring at a rate slower than species accumulation (**Figure S3F-G**). The *frequency* morphospace was explored to a greater degree (37%; **Figure S3E-G**) that evolved in tandem with species accumulation until an increase in variation emerged ~1.5 Ma (**Figure S3F-G**). Finally, while the *transitions* morphospace was slightly less explored than *frequency* (30% covered; **Figure S3F**) it showed multiple pulses of diversification that outpaced species accumulation (**Figure S3E-F**) and filled earlier than the other traits (**Figure S3G**).

To what extent might specific components of these traits – movements or the transitions between them – be evolving at different rates to account for these patterns? Strikingly, we found that frequencies of specific movements did not evolve uniformly but rather varied >4 fold across behavior space with especially rapid rates in regions associated with high translational velocity, turning at high speed, and stopping (**Figure 4A**). Similarly, transitions between specific movements varied in rate disproportionately (**Figure 4B**). The recurrence of high translational velocity and fast turning movements, and the transitions between them, evolved most rapidly (**Figure 4B**). Furthermore, comparing species based on overall variation in these patterns indicated evolutionary convergence. A phylogeny produced from variation in transitions regrouped *D. virilis, D. arizonae,* and *D. mauritiana* with a subset of the *D. melanogaster* clade and strains (**Figure 4C**). A similar result was evident for the frequency of movements (**Figure S4A**), both of which reflected the species groupings present in their respective morphospaces (**Figure 3C**, **S3A**).

**Figure 4:**
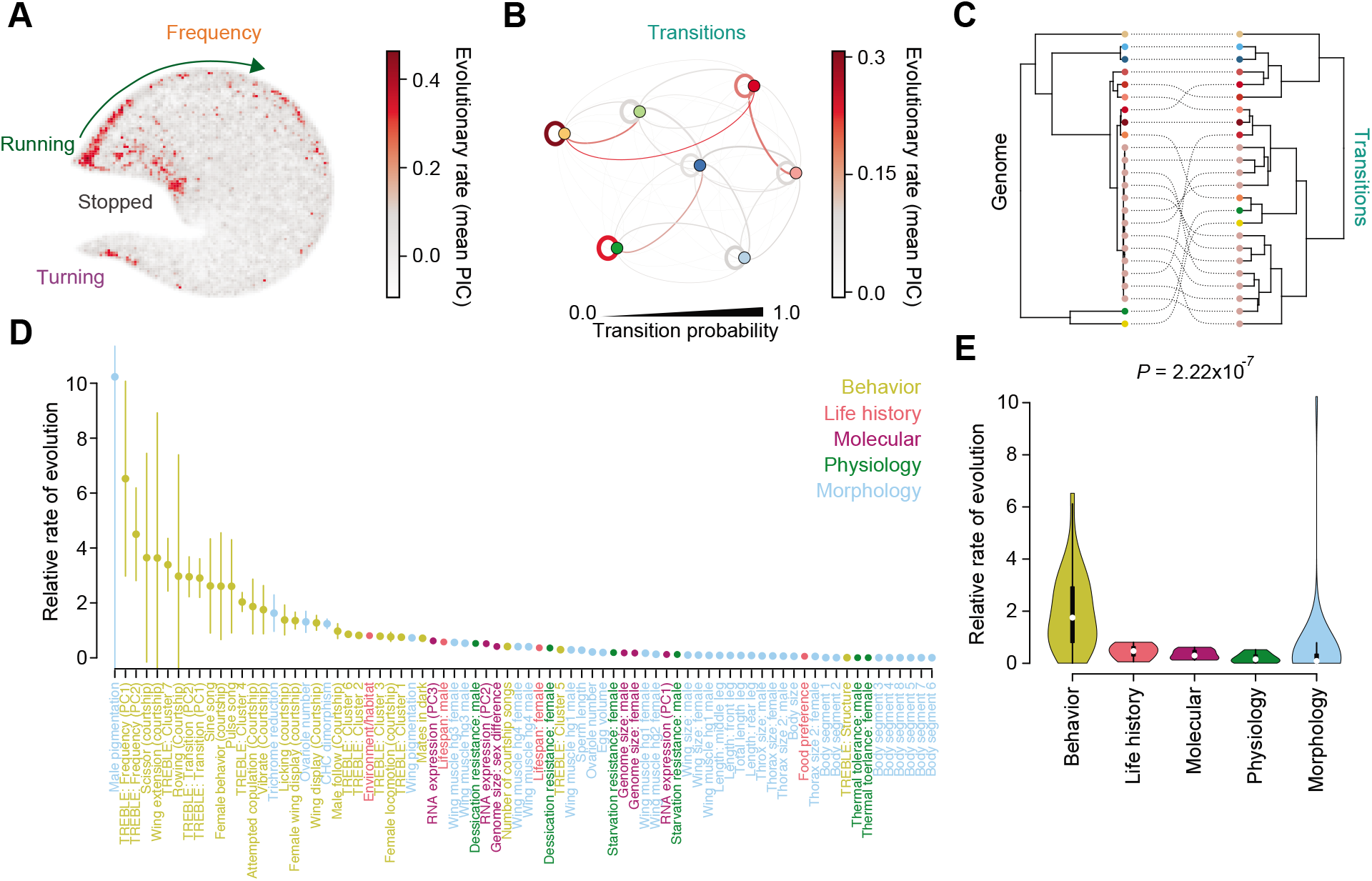
Rapid, specific, and convergent behavioral evolution in *Drosophila*. **(A)** The evolutionary landscape of*frequency.* Darker colors correspond to greater evolutionary rates (measured by phylogenetic independent contrasts; PIC). **(B)** The evolutionary landscape of *transitions.* Transition probability is represented by the thickness of the lines connecting nodes (corresponding to Louvain clusters; see Methods). Evolutionary rate corresponds to darkness of color (phylogenetic independent contrasts; PIC). **(C)** Cophyloplot comparing the fossil-calibrated whole-genome phylogeny (left) to a phylogeny made from variation in transitions (right). Species are represented by colored nodes at the terminal tips of both phylogenies, with their corresponding positions indicated by dotted lines. **(D)** Distribution of relative rate of evolution across five categories of *Drosophila* traits: behavior, life history, molecular, physiology, and morphology. Points correspond to median values, bars represent standard error. **(E)** Violin plot of relative rates of evolution given *Drosophila* trait type (Kruskal-Wallis test *P*-value).

Taken together, these results imply a hierarchy in the evolution of drosophilid locomotion: *transitions* are most malleable, *frequency* evolves slightly less rapidly but with greater variance than transitions, and *structure* is largely conserved. We note that recent work described a similar relationship between *frequency* and *structure* using different methods and a smaller set of species, but could not resolve *transitions* (Hernández et al. 2021). Furthermore, these findings suggest that the rapid evolution of *frequency* and *transitions* resulted in convergence of the walking repertoires of multiple species and strains. Finally, these findings reveal that walking is organized around two fundamentally different strategies, one that is built on longer sequences of high velocity runs and turns and the other on shorter sequences of lower-velocity movements.

### Behavior evolves more rapidly than other traits

Given the hierarchical and rapid evolution described above, we wondered how the diversification of walking might compare to that of other traits. To do so, we conducted a meta-analysis of the evolution of 78 behavioral, morphological, molecular, physiological, and life history traits (**Table S2**). We normalized trait measurements and used the fossil-calibrated genome tree to calculate evolutionary rate and phylogenetic signal for each (Methods). For these traits, the relative rate of evolutionary change varied over an order of magnitude (**Figure 4D**). Strikingly, of the 25 most conserved traits, only one was behavioral and, as expected, was *structure.* Of the 25 most rapidly evolving traits, 21 were behavioral including aspects of *frequency*, *transitions*, and a variety of courtship behaviors. Importantly, the remaining 4 rapidly evolving traits were morphological and tightly associated with courtship which is known to evolve rapidly in drosophilids (Spieth 1952, O’Grady & Markow 2012, Jezovit et al. 2017). Reflecting this pattern, behavioral traits showed the greatest rates of evolution (**Figure 4E**).

## Discussion

Here we dissected the evolution of walking in a phylogenetically diverse clade, fruit flies of the genus *Drosophila.* A universal behavior space captured the full range of drosophilid walking kinematics. Variation in hierarchical aspects of the walking behavior space – *structure, frequency,* and *transitions* – was captured by individual morphospaces, each with a unique tempo and pattern of evolutionary diversification. While *structure* was conserved, *frequency* and *transitions* displayed multiple pulses of phenotypic evolution, resulting in species and strains that converge on common movement patterns. Furthermore, behaviors were associated with significantly faster rates of evolution than morphology, life history, physiology, and molecular traits, results that are consistent with previous studies (Gittleman et al. 1996, Blomberg et al. 2003). Whether this pattern holds across other systems will be of great interest. Such comparative work could resolve the long-standing debate of whether behavior is a facilitator, or inhibitor, of phenotypic variation during organismal evolution (Larmarck 1809, Darwin 1859, Huxley 1942, Mayr 1982, Wcislo 1989, Blomberg et al. 2003). In particular, notwithstanding the need to perform comparable analyses using other behaviors and clades, by revealing relatively rapid evolutionary change in the behaviors of one model clade, our work supports the idea that behavior can indeed facilitate the emergence of phenotypic variation.

Our data demonstrate that sequences of motor actions, rather than individual movements, have evolved rapidly. Furthermore, divergent species and strains have converged on similar behavioral patterns multiple times, while closely related strains that live in comparable natural environments can be highly divergent behaviorally. Thus, the evolutionary patterns we observed appear robust to the specifics of our behavioral assay. In some cases, behavioral convergence has occurred independent of life history and ecological background. For example, species from tropical (*D. mauritiana*) and temperate (*D. virilis*) habitats have converged on similar behavioral repertoires to those seen in the cosmopolitan *D. melanogaster*. In the same vein, very closely related *D. melanogaster* strains isolated from a single sub-region in Africa can also diverge markedly in their walking behavioral repertoires, spanning distances in behavior space comparable to that seen in divergent species pairs. Future work should explore the evolutionary and co-evolutionary relationships between behavioral structure and traits such as physiological regulation and anatomical differences across *Drosophila* species.

Studies of a wide range of both social and non-social behaviors, in many animals, have identified discrete, modular elements that can be arranged in temporal sequences of varying flexibility (Tinbergen 1950, Tinbergen 1951, Dawkins 1976, Herrel et al. 2001, Glaze & Troyer 2006, Xu et al. 2012, Berman et al. 2014, Jin & Costa 2015, Gomez-Marin et al. 2016, Katsov et al. 2017, Wiltschko et al. 2015, Duistermars et al. 2018, Marques et al. 2018, Tao et al. 2019, Kaplan et al. 2020, Johnson et al. 2020). Here we compare the structure of a common behavioral repertoire across a densely sampled set of species and strains to show that the apparent focus of rapid evolutionary change lies in changing the temporal sequences of individual movements, with substantially less variation in the fine structure of the movements themselves. Thus, the modularity of behavior seen in individual species in fact appears to reflect the structure of evolutionary change. These observations, in tandem with comparisons to other traits, suggest that the diversification of walking arises first via changes in neural control, as opposed to biomechanical or morphological mechanisms that would alter the fine structure of individual movements. In this case, behavioral diversification may be tightly coupled with neural evolution and labile over even very short evolutionary timescales, such as those seen within a species. Thus, the architecture and function of neural circuits appears to both enable complex behavior and facilitate its rapid diversification.

## Acknowledgements

We thank members of the Clandinin, Tuthill, and Matute labs and members of the NIH U19 FlyLoops consortium for useful discussion. This work was supported by the NIH (U19NS104655: T.R.C., J.C.T.; R01GM121750: D.R.M.), the Simons Foundation (T.R.C.), the Stanford School of Medicine (R.A.Y.), the New York Stem Cell Foundation (J.C.T.), a Searle Scholar Award (J.C.T.), a Klingenstein-Simons Fellowship (J.C.T.), a Pew Biomedical Scholar Award (J.C.T.), a Sloan Research Fellowship (J.C.T.), and NSF Dimensions of Biodiversity award 1737752 (D.R.M.). J.C.T. is a New York Stem Cell Foundation – Robertson Investigator.

Lastly, we want to acknowledge that our laboratory at Stanford University stands on the ancestral lands of the Muwekma Ohlone Tribe (http://muwekma.org/home.html).

## Author Contributions

R.A.Y. and T.R.C. conceived the study. L.B., S.H., A.K., A.S., E.S., and R.A.Y. developed the Coliseum and FlyVR software. R.A.Y. collected the walking data set. S.L., B.P., and J.C.T. performed gait tracking. J.C., A.S., and D.R.M. compiled and prepared the whole-genome phylogeny. R.A.Y. curated the data and performed statistical analysis. R.A.Y. and T.R.C. prepared the manuscript.

## Materials & Correspondence

Correspondence and material requests should be addressed to.

## Materials and Methods

### Fly strains

All stocks were kept at 25°C on molasses-based food and reared under a light-dark cycle of 10:10 h. The following stocks were obtained from National Drosophila Species Stock Center: *D. willistoni* ((Heed) H57.30), *D. santomea* ((Gompel)STO.4), *D. persimilis* (2529.6), *D. mauritiana* (E18912 MS17), *D. virilis* (3367.1), *D. arizonae, D. yakuba, D. sechellia, D. simulans, D. pseudoobscura, D. erecta,* and *D. teissieri.* Strains of *D. melanogaster* were collected in Southern Africa, the ancestral range of the species (Coughlan et al. 2021).

### Behavior

#### Apparatus

The Coliseum is an enclosed 1m x 1m arena for measuring the unconstrained walking of fruit flies. The arena is sealed from external light via Velcro-attached curtains on the sides and solid walls on the top and bottom. Flies are released into the arena through a hole in the floor by means of an automated dispenser consisting of a vial filled with flies and a servo-gated exit. Flies enter the arena by crawling up the exit channel. An optical sensor detects that a fly has entered the channel and immediately closes the gate behind the fly to both avoid releasing multiple flies simultaneously and to prevent flies from returning to the vial from the arena. Once a fly is in the arena, it is edge-lit by IR LEDs around the perimeter of the floor and recorded from above by a high-definition camera outfitted with a zoom lens that is sufficiently powerful to capture anatomical details at high resolution. To keep the fly in its field of view the camera is mounted on a 2-axis CNC mill (grbl with gShield + Arduino UNO; stepper motors are SM42HT47-1684B) that updates the camera position as the fly moves.

The position of the CNC mill is controlled by the software flyvr (github.com/ClandininLab/flyvr) by tracking the position of the fly and the camera simultaneously. The fly’s position in the camera is first computed by thresholding and extracting the pixels representing the fly, identifying the head-tail axis and orientation, and then calculating the in-frame coordinates of the fly’s center. This relative position is then summed with the position of the camera to calculate an absolute position in the Coliseum which is used to update the stepper-motor coordinates of the CNC and thus keep the camera in sync with the fly’s movement. The absolute position and heading angle of the fly are recorded for each frame (at 100Hz) and outputted by flyvr at the end of each trial.

#### Behavior experiments

Immediately after eclosion, virgin female flies were sorted into vials of 10-20 flies and reared in light-dark chambers (12:12h) at 25°C. 2-4 day old flies were used for tracking in the Coliseum. On the day of the experiment, individual vials would be loaded directly into the automated dispenser and flies would enter the Coliseum one at a time. Flies were allowed to explore the arena freely for up to 20 minutes, after which the animal would be manually removed from the chamber. The floor of the Coliseum was cleaned with 70% ethanol between trials to remove odorants or other stimuli that may affect patterns of locomotor behavior. All experiments were conducted during the same time window to align with the light-dark cycle, from roughly CT0-CT3.

Age, diet, circadian rhythm, and environment were controlled to facilitate interspecific comparisons, potentially influencing individual species’ expression of locomotor behavior. However, a standardized diet has minimal effects on the behavior of diverse species, even on specialist species such as *D. sechellia, D. arizonae,* and *D. erecta* (Shultzaberger et al. 2019). Future work comparing the locomotion of wild-caught flies in natural conditions would be useful for corroborating the controlled laboratory experiment presented here.

### Generating a universal behavior space with TREBLE

#### Determining window size

Following previous work on the kinematics of walking in *D. melanogaster* (Katsov et al. 2017, York et al. 2021) we calculated the following kinematic parameters for each trial: translational velocity (cm/second), angular velocity (degree/second), and sideslip (cm/second). Per-frame position and heading angles from the Coliseum were used for these calculations. As in York et al. 2021, angular and sideways velocity values were normalized to the first frame in the window and proceeding velocity values were adjusted to ensure that the second frame was always positive. Since we weren’t concerned with turning direction the absolute value of angular velocity was used. To control for rare occasions of jitter in the camera movement we smoothed each velocity parameter using a Nadaraya-Watson kernel estimate (ksmooth function in R; bandwidth = 0.25; R Core Team 2020) and down sampled the data to 30Hz (which removed noise without affecting the structure of the velocity parameters).

A core component of the TREBLE framework is identifying the temporal structure of a behavioral dataset by iteratively sampling data in increasing window sizes and empirically analyzing variation in the downstream behavior spaces (York et al. 2021). Here, we performed this iterative window search by randomly sampling 50 trials and testing the effect of window sizes (from 10ms to 1 second, 20ms intervals) on the structure and temporal properties of behavior space (**Figure S1E-I**). The goal of this procedure is to identify a window size that minimizes variation across replicates in both the structure and temporal sequencing of movement through behavior space. Variation in behavior space structure was assessed by measuring the Procrustes and mean inter-point Euclidean distances between all replicates (**Figure S1F-G**). Here, Procrustes distance is a measure of the global differences between two spaces (York et al. 2021) while mean inter-point Euclidean distance provides a local measurement (i.e. between adjacent points) of structural variation. In both cases the mean stabilized, and variance decreased, at windows around 300ms in size.

Temporal variation was measured by analyzing the timing of recurrence in behavior space, based on the concept that these are essentially state spaces reflecting the function of a continuous dynamic system (York et al. 2021). Recurrence was measured by calculating the duration of time it took to return to the local neighborhood of each point in behavior space. The proportions of points that were recurrent as a function of 10ms temporal bins, ranging from 0 to 2 seconds, were then measured (**Figure S1E**). From this we obtained the mean recurrence time across all temporal bins (**Figure S1H**) and the overall proportion of recurrent points across all bins (**Figure S1I**) for each window size. As has been seen previously in studies of fruit fly walking (York et al. 2021), there was a narrow range of window sizes (~140-360ms) that displayed a peak of recurrence at around 250ms (**Figure S1E**) and dissipated with increasing window size. Taken together, these analyses indicated that a window size of 333ms optimally reduced variation in the structure of behavior space while capturing the temporal patterns present.

#### Embedding temporal windows into behavior space

We next produced windows for all trials using the size identified above. For each frame of a given trial (denoted here as frame i), and for a window size w, we individually extracted the kinematic parameters (x) for time i:i+w and then concatenated them linearly into a single vector of length 3x. These window vectors were then collected into a single matrix spanning all flies and containing 3,890,645 windows. Further details of this procedure can be found in York et al. 2021. We then used the R implementation of the UMAP algorithm (McInnes et al. 2018) to non-linearly embed the windows into a 2-dimensional behavior space containing all flies from all species and strains. To facilitate downstream analyses, we also produced version of the space with simplified 2-d coordinates by decomposing each point onto a grid of size *n* bins x *n* bins using the TREBLE function bin_space. Most analyses described below use a 64×64 grid size, containing 4,096 unique bins, unless otherwise noted.

#### Characterizing behavior space

Stereotypy in movement through behavior space was assessed by creating a vector field representation. Using the 64×64 grid representation of behavior space, we identified all points associated with a given bin and then calculated the mean direction for all instances of a trajectory leaving that bin (in xy coordinates). Mean direction was represented by plotting the mean trajectory out of each bin via an arrow, the direction and magnitude of corresponded to the xy coordinates calculate above (as in **Figure 2D**). Arrow color reflected the polar coordinates of the vector for each bin. To measure regional covariation (as in **Figure 2E**), we first identified the 20 nearest neighbors of each bin in the vector field representation (64 × 64 grid) using the function get.knn from the R package FNN. We then calculated the covariation of each bin’s vector with its 20 nearest neighbors, reasoning that regions with similar movement patterns would show increased covariation across neighboring bins.

Variation in the percent of behavior space covered by individuals and species/strains (**Figure 2J**) was measured by calculating the number of unique bins in behavior space were visited as a function of the total number of bins. The significance of per-bin variance as function of species (**Figure 2K**) was measured using a Kruskal-Wallis test. First, we calculated the total proportion of time spent in each bin (as a fraction of overall trial time) for all flies. We then ran a Kruskal-Wallis test (kruskal.test function in the R package stats) for each bin, comparing species and using the bin-wise proportions from each fly of a given species to account for intra-specific variation (R Core Team 2020). The resulting *P*-values were adjusted for multiple tests using Bonferroni correction.

Patterns of temporal sequencing through behavior space were inferred by calculating the autocorrelation of position in behavior space. To do so, we assigned each bin in behavior space a unique 1-d identifier, creating a single numeric representation of behavior space position (a metric outputted by bin_space). This vector was then used to calculate autocorrelation using the acf function in R (lag time = 3 seconds), for all flies (**Figure 2L**) and for all species (**Figure 2M**).

#### Behavior space power tests and intra-inter-species variance

Animal behaviors can be influenced by a variety of factors – such as context, physiological state, or personality and, in addition, genotype (Wcliso 1989). If the variance of a given behavior is strongly driven by contextual factors, then identifying consistent species differences may be difficult or impossible. With this in mind, we sought to calculate the intra-individual variability in behavior space statistics and to compare this to the variance observed with and between strains and species.

To address the first goal, we performed a power test exploring the number of individuals needed to capture the overall structural and frequency statistics within behavior space for each species. First, we calculated probability density functions (pdfs) of xy-coordinates in behavior space for each species using all individuals (grid points = 100, bandwidth = 1). We then compared pdfs calculated using a range of included trials (1 to 10 trials) to the full pdf for each species using correlation. To factor in intra-trial variability, we bootstrapped each trial sample size 10 times (i.e., 2 trials were randomly chosen 10 times and correlated to the full pdf; then 3 trials were randomly chosen 10 times...), randomly permuting which trials were used per species. Comparing mean correlations across permutations, we found that each species and strain had a high level of autocorrelation with a small number of trials included, for most reaching a correlation >0.9 using just 6 trials and converging >0.95 with 10 trials used (**Figure S2A-B**). These results indicated that, for the given behavior and experimental set up, intra-species variation in walking could be captured well within the sample sizes collected (**Figure S2A**).

We next assessed the relationships of intra-individual, intra-species, and inter-species variation by bootstrapping behavior and comparing distributions in behavior space using cosine similarity. We first selected trials longer than 5,000 frames (166.66 seconds), resulting in 373 trials. Intra-individual variance was measured by randomly selecting 2,000 time points 10 times per trial (without replacement), resulting in a total 3,730 permutations. A pdf (grid points = 100, bandwidth = 1) was calculated each permutation. The intra-trial similarity was calculated by computing the cosine similarity of the pdfs from the 10 permutations per trial (**Figure S2B**). Intra-strain variance was computed in the same fashion except for 100 bootstraps we performed per strain (**Figure S2B**). Inter-specific variance was measured by bootstrapping xy coordinates (2,000 per shuffled) from the full data set 1,000 times (**Figure S2B**). The distributions of all three measures were compared using a Kruskal-Wallis test followed by a post-hoc Dunn’s test (dunn.test function in the R package dunn.test). We found that all three significantly differed. Intra-individual similarity was greater than intra-strain while intra-strain similarity was greater than interstrain (**Figure S2B**).

### Gait analysis

#### Limb tracking

Behavior videos were first contrast enhanced using custom Matlab scripts to optimize the view of the animal’s legs. The head, thorax, abdomen, and leg tarsi were automatically tracked from the top-down videos using DeepLabCut and Anipose (Mathis et al. 2018, Karashchuk et al. 2020). Tracked data points were used to analyze walking kinematics using a custom Python script (https://github.com/Prattbuw/CODE-Evolution-of-Drosophila-Walking). Position time series data associated with the tarsi was smoothed using a moving average with a time window of ~80ms. Head, abdomen, and leg tarsi positions were normalized to the thorax to calculate positions relative to the body. Heading angle was calculated based on head and thorax positions relative to an allocentric reference (i.e. experimental chamber). Tarsi positions were rotated and translated based on deviations from a common heading angle (allocentric angle of 0°). This transformation was necessary for extracting stance and swing onsets as this maximizes the position signal of each leg tarsi signal along the body axis. Steps were extracted by identifying the peaks and troughs of the tarsi position signals. Steps consist of two phases: swing (trough to peak) and stance (peak to trough). Number of legs in stance was calculated by summing the number of legs touching the ground at each sample point. Leg phase (as in **Figure 2F**) was calculated using a Hilbert transformation of the position of each leg at all frames (HilbertTransform function in the R package hht (Bowman et al. 2013)). The output values were then smoothed using a Savitzky-Golay filter (order = 3, filter length = 15) using the function sgolayfilt in the R package signal (Signal developers 2014).

#### Gait analysis

Flies were considered walking when body velocity was greater than 5 mm/s, the maximum likelihood of each of the six tracked points was greater than 0.99. We identified a small number of rare cases in which flies were estimated to have 0 or 1 legs in stance while moving at low velocities. Given this, all instances in which these stances were detected at body velocities less than 10mm/s were replaced with 6 leg stances. We then identified all continuous walking bouts longer than 333ms, yielding 1,123 bouts representing 216,729 time points.

The walking bout data set was then used to train Hidden Markov Models (HMMs) modelling the states underlying the number of legs in stance over time using the R package depmixS4 (Visser 2010). We used the function depmix to train three models (2 hidden states, 3 hidden states, 4 hidden states) using the formula stance ~ 1. We then optimized the model parameters using expectation maximization via the fit function in depmixS4. The fits of the three models were compared using log likelihood AIC, and BIC (**Table S1**), revealing that a model with 3 hidden states fit the data best and paralleling previous observations in fruit flies (Mendes et al. 2013, Wosnitza et al. 2013, Isakov et al. 2016, Pereira et al. 2019). Per-frame state designations were estimated from the posterior probabilities of the models. The distributions of HMM states in behavior space were calculated by identifying the xy coordinates in behavior space for all time points in which a given state occurred. These were then used to calculate probability density functions as function of behavior space position for each state (as in **Figure 2I**) using the function kde2d in the R package MASS (grid points = 200, bandwidth = 2).

### Phylogenetic analyses

#### Calculating structure, frequency, and transitions

*Structure* was measured by identifying the unique bins that each species or individual (depending on the analysis) visited. A 64 × 64 binary matrix was generated and bins visited were filled with 1 while bins not visited were filled with 0.

*Frequency* was inferred by calculating probability density functions (pdfs) from xy position in behavior space. Here, greater density in a specific region would reflect a higher frequency of occurrence of the movement represented by that portion of behavior space. We calculated pdfs for all individuals using the function kde2d in the R package MASS (grid points = 100, bandwidth = 1). To enable comparison each pdf was normalized by dividing all values by the maximum. From these we calculated a mean pdf for each species by averaging each point across all individuals of a species, in addition to the standard error for each bin (used for downstream phylogenetic analyses).

*Transitions* were identified by first clustering points in behavior space based on their graph properties. Many clustering methods identify structures based on the density of points across some number of dimensions. However, the TREBLE behavior space contains information about both point density *and* the temporal sequencing between points. We therefore sought a clustering method that could capture both important aspects. Furthermore, to facilitate rapid comparisons across individuals and species, we prioritized methods that minimized assumptions and extensive model fitting. We opted to use Louvain clustering, a graph-based approach common in other non-linear dimensionality reduction application such as scRNA-seq (Levine et al. 2015). First, we created an undirected graph with two columns using the function graph_from_data_frame in the R package igraph in which the first column represented the xy coordinate *from* which a trajectory was leaving while the second was the xy coordinate the trajectory was going *to.* This resulted in a graph with 3,890,645 rows corresponding to a full set of feature windows and their corresponding 2-d movement patterns in behavior space. We then used the igraph function cluster_louvain to do the clustering. The procedure yielded 7 clusters that corresponded to recognizable features based on point density and movement through behavior space (**Figure S2C-E**). Furthermore, each cluster was associated with a characteristic joint distribution of kinematic parameters used (**Figure S2B**), reflecting distinct components of movement. Transition probabilities between Louvain clusters were inferred using Markov models. We created models for all flies using the function markovchainfit in the R package markovchain (Spedicato 2017) using the Louvain cluster designations from cluster_louvain as input. The transition matrices for each fly were extracted and averaged per-species to create a mean transition matrix for each. The standard error of each transition was also calculated per-species for downstream phylogenetic analyses.

We calculated morphospaces for *structure, frequency,* and *transitions* using PCA (**Figure 3C**, **S2C-E**). The input matrices for PCA were produced by linearizing and combining the species mean values for each trait. For example, the 64 × 64 *frequency* matrices were linearized into 4,096 element vectors and horizontally combined to create a 4,096 × 24 matrix that was then used to run a PCA.

#### Mapping trait evolution

Phylogenetic signal was measured via Blomberg’s *K* using the R package phytools (Revell 2012). For the traits *structure* and *frequency,* we calculated phylogenetic signal for all bins in behavior space, treating each as a unique phenotypic measurement, and using the species means and standard errors calculated above. We assessed the extent to which the number of bins chosen for these traits may affect the calculation of phylogenetic signal by calculating the phylogenetic signal distributions of a range of resolutions (2×2 to 100×100) for the trait *frequency.* Comparing mean phylogenetic signal revealed that the measure began to level-off around a resolution of 20×20 bins and become stable at a resolution of 30×30 bins (**Figure S2F-G**), demonstrating that a relatively broad range of possible resolutions could be used in comparing this trait across species. For *transitions* we calculated phylogenetic signal of each transition probability contained in the mean species-level transition matrices, also factoring in the standard error. This resulted in distributions of phylogenetic signal across all dimensions of the three traits (**Figure 3D**). Variation in these distributions was tested using a Kruskal-Wallis test followed by a post-hoc Dunn’s test.

To compare the temporal patterns of trait evolution among *structure*, *frequency*, and *transitions* we computed variable rates models using the R implementation of the software BayesTraits (evolution.rdg.ac.uk, v.3), and its wrapper btw (github.com/rgriff23/btw). To facilitate multivariate comparisons, we used independent contrasts models for all three traits and applied Markov-Chain Monte Carlo chains run with 10,010,000 iterations and thinned every 1,000. For *frequency* and *transitions* we modelled all PCs that accounted for >90% of variance in the data (13 and 4, respectively). The first two PCs were used for *transitions.* Three models were created for each trait. We tested for convergence of all models using the Gelman-Rubin diagnostic in the R package coda (function gelman.diag, Plummer et al. 2006; 3 chains), requiring a value of 1 to proceed with analysis. We required models to have effective sizes of at least 200 and selected the best model for each trait via Bayes factor using btw. To compare the distributions of relative evolutionary rates, we extracted the rates reported at each node in the tree by BayesTraits and normalized them by the maximum value for each trait (as seen in **Figure 3E**). Variation in rate distributions was tested using a Kruskal-Wallis test followed by a post-hoc Dunn’s test.

We performed an adapted version of the analyses in Cooney et al. 2017 and Ronco et al. 2021 to estimate morphospace filling over time. First, we estimated ancestral states for each trait using the fastAnc function in phytools using the mean rate-transformed tree from BayesTraits. We then calculated the values of each trait along the species tree in time intervals of 0.1 million years. At each interval, trait values for all extant branches were identified and then linearly predicted between nodes. To evaluate morphospace filling we performed this procedure for the first two PCs of each trait, as seen in **Figure S3E**. To estimate the percent of morphospace filled as a function of time (**Figure S3F**), we calculated the cumulative coverage of the xy coordinates of the 2-d morphospaces in 0.1 million year intervals.

The evolutionary rates of specific components of *frequency* and *transitions* were inferred by computing phylogenetically independent contrasts (PIC) with the R package ape (pic function; Paradis & Schliep 2019). For *frequency*, PIC was calculated for every bin in behavior space. For *transitions*, PIC was calculated for each transition probability between Louvain clusters.

#### Trait meta-analysis

To compare the patterns observed here to other *Drosophila* traits, we conducted a review of the fruit fly evolutionary literature and identified studies that measured traits in at least 7 species present in our phylogenetic tree. This yielded measurements for 56 traits across 18 individual studies (see **Table S2** for citations). In addition, we incorporated *frequency* (PC1, PC2), *transitions* (PC1, PC2), *structure* (% of behavior space covered), and the frequency of occurrence in the 7 Louvain clusters. Overall, the final data set contained 78 traits with 105 species represented at least once (**Table S2**; **Supplementary data S1**).

To facilitate comparison across traits, each measure was converted to a 0-1 scale by first adding the absolute value of the minimum and then dividing by the maximum value. We then inferred evolutionary rate (σ^2^) using a single-rate Brownian motion model in phytools. Rates were then compared across trait types using a Kruskal-Wallis test followed by a post-hoc Dunn’s test.

Estimates of evolutionary rate can be biased by sample size and the phylogenetic distance of the clade being compared *(65).* To test if such biases were present in our data set, we created a linear model predicting σ^2^ from sample size and distance (inferred by the maximum branch length of the phylogenetic tree for a given subset of species; lm function in stats). While sample size and distance accounted for very little of the variation in σ^2^ (R^2^ = 0.07), and sample size did not significantly predict the outcome variable (*P* = 0.82), we did find that distance was marginally significantly predictive (*P =* 0.02). If this association between phylogenetic distance and evolutionary rate were to be unevenly distributed across trait categories, then the observed rate differences might have arisen from artifacts rather than real signal. To control for this potential limitation, we compared the residuals of the linear model (i.e. σ^2^ with the effects of sample size and distance regressed out) using a Kruskal-Wallis test. We found that the five trait types still differed significantly (*P* = 8.7 × 10^-5^) (**Figure S4B**).

---

**Figure S1:**
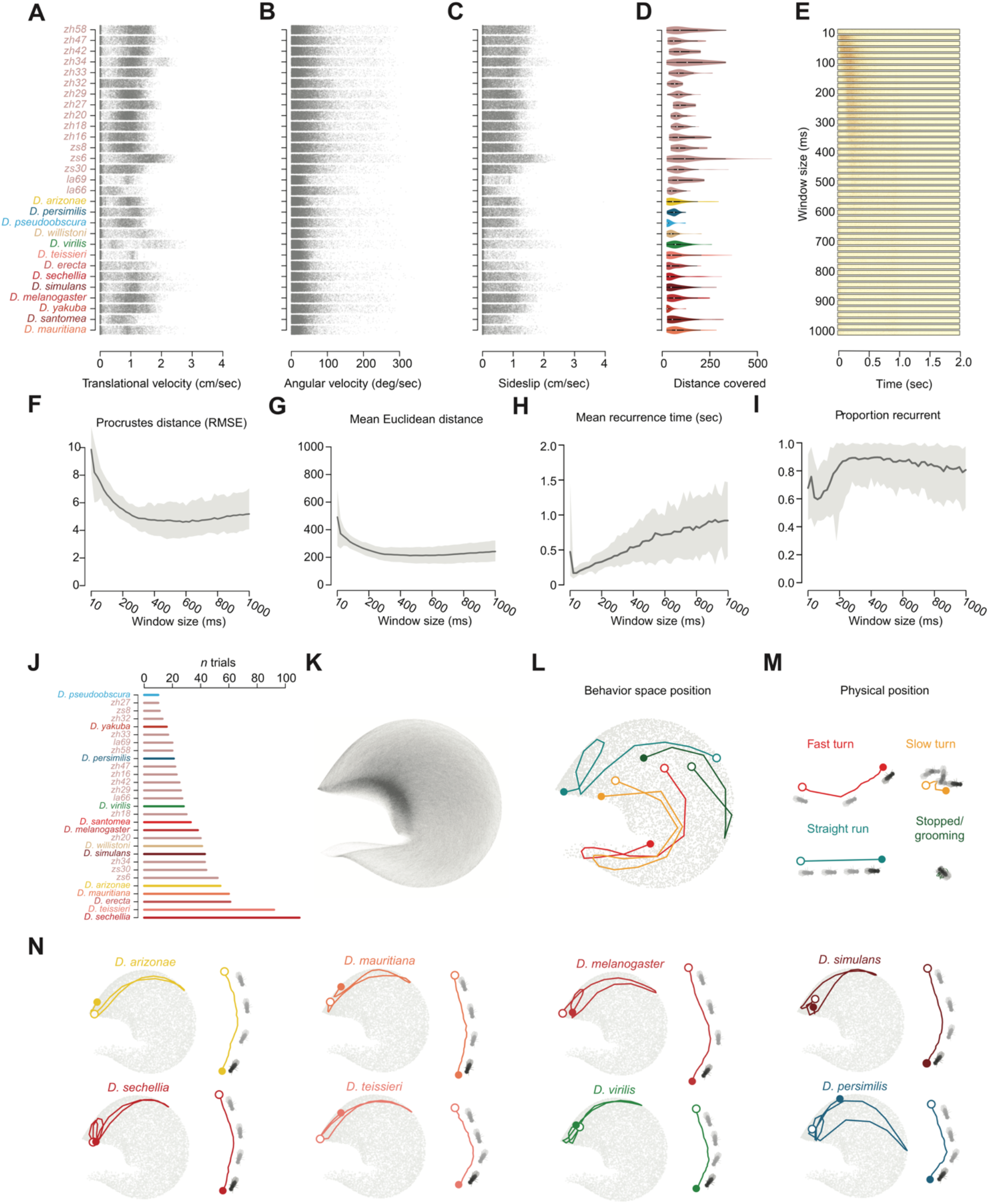
Characterizing the walking data set and TREBLE embedding. **(A-C)** The distributions of absolute angular velocity (**A**), translational velocity (**B**), and absolute sideslip **(C)** calculated from all flies of each genotype. **(D)** Violin plot of the distances covered per genotype. **(E)** Recurrence plot testing the effect of window size on behavior space for the 50 trials tested. Window size on the y-axis, the time delay after visiting a given point is on the x-axis (Methods). **(F-G)** The distributions of Proscrustes distance measures (RMSE) (**F**) and mean inter-point Euclidean distance (**G**) as a function of window size calculating during the iterative window search procedure in TREBLE. **(H)** Mean recurrence time (seconds) as a function of window size. **(I)** Total proportion of points that are recurrent as a function of window size. Each measure (**F-I**) was calculated over the 50 replicate behavior spaces (i.e. n = 50 per window size). Darker line is the median value, lighter gray represents standard error. **(J)** Barplot of the number of trials per species and strain used to create the walking behavior space. **(K)** Continuous pathways through behavior space. All temporally continuous points in behavior space are connected by lines, highlighting common pathways and areas of greater density. **(L-M)** Example behaviors in physical space (**D**) and their corresponding positions in behavior space (**C**). **(N)** Examples of the same behavior across 8 exemplary species. For each species, positions in behavior space are plotted on the left and movements in physical space are on the right.

**Figure S2:**
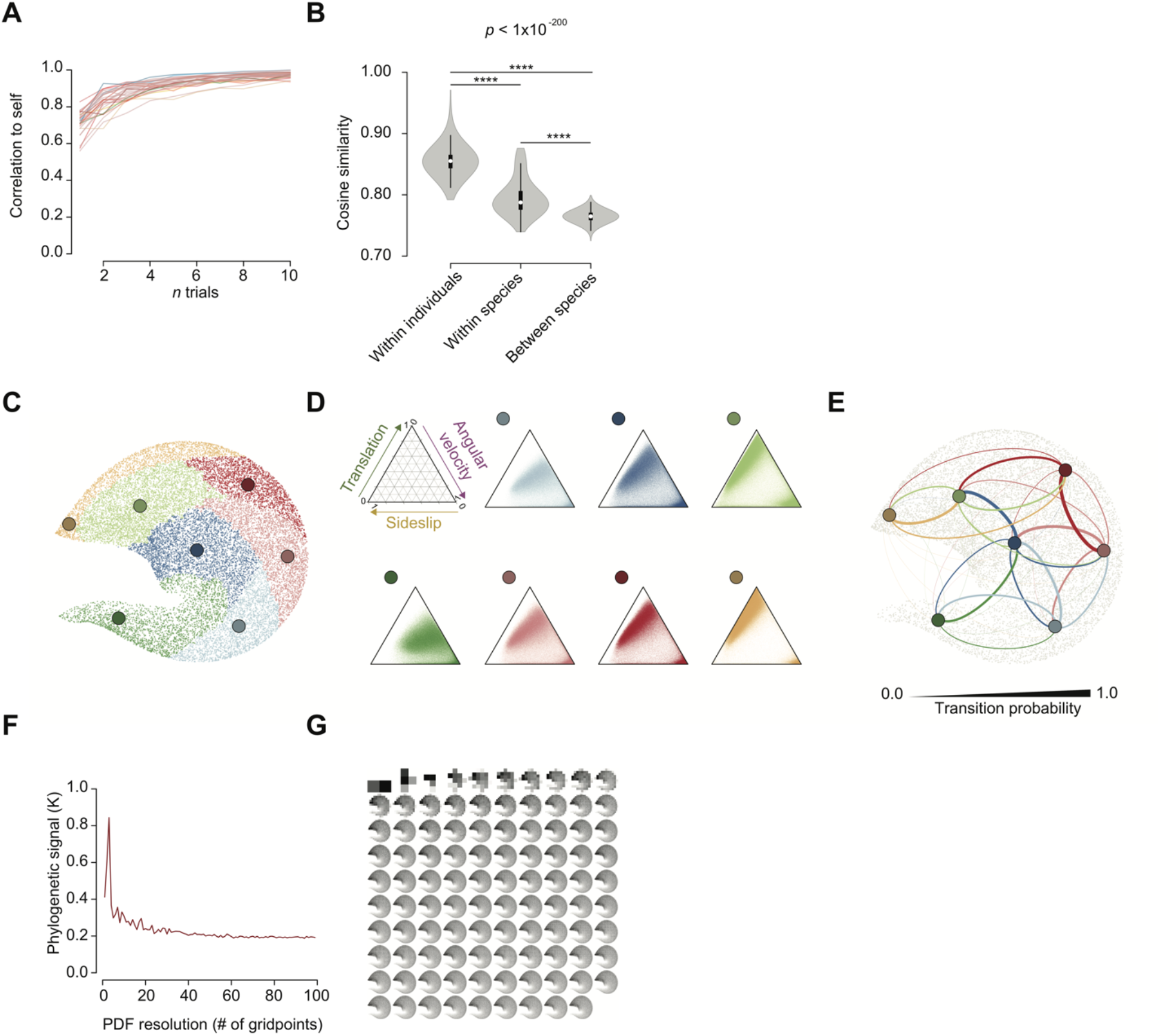
Statistical and evolutionary properties of the universal behavior space. **(A)** Bootstrapped correlation values (means) comparing the behavior space distribution of randomly chosen subsets of trials to the overall distribution of a species or strain. Colored by species and strain as in **Figure 1C**. **(B)** Violin plots comparing cosine similarity distributions of randomly sampled behavior space positions within individuals and species and between species. **** *P*<0.0001. Kruskal-Wallis test followed by Dunn’s test. **(C)** The distribution of the 7 identified Louvain clusters, colored within the boundaries of each. **(D)** Ternary plots displaying the joint distributions of translational velocity (‘Translation’), angular velocity, and sideslip for each of the 7 Louvain clusters. Values of each kinematic parameter increase with the direction of the arrows in the first panel. Each ternary plot contains all points that are embedded in the behavior space. **(E)** Transitions between Louvain clusters. Clusters are represented by colored points arranged in the same positions as **(C)**. Transition probabilities are represented by the thickness of the lines connecting each point (and correspond to the legend below the panel). **(F)** The distribution of phylogenetic signal (Blomberg’s *K)* as a function of probability density function resolution for *frequency.* For each resolution, phylogenetic signal was calculated as in **Figure 2D**. Resolutions of 2-100 gridpoints (interval of 1) were tested. **(G)** The 2d probability density functions used for calculating phylogenetic signal in **a**, ranging from 2-100 gridpoints.

**Figure S3:**
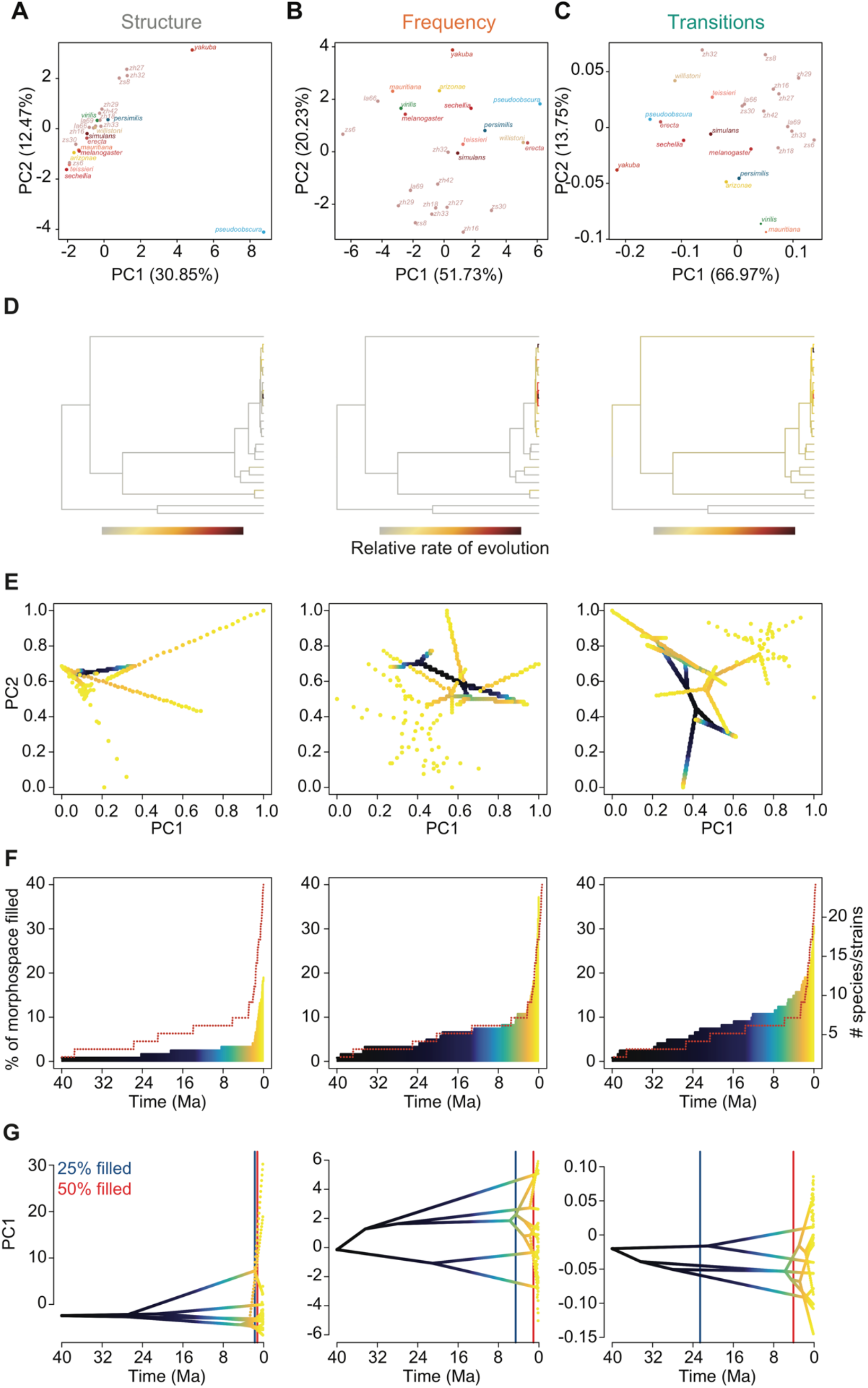
Morphospace filling. **(A-C)** Morphospaces for *structure* (**A**), *frequency* (**B**), and *transitions* (**C**). Morphospaces were created from the first 2 principal components of a PCA run on each trait (see Methods). **(D)** Species trees with relative rate of evolution indicated by color. **(E)** Movement through morphospace over time. Time (Ma) is indicated by color (corresponding to **E**, **F**, and **G**). **(F)** Proportion of extant morphospace filled as a function of time (calculated from the analyses in **E**; see Methods). Species accumulation is represented by the red dotted line. **(G)** Morphospace densities over time for PC1 of each trait. Time points for 25% and 50% of morphospace filled are indicated by blue and red lines, respectively.

**Figure S4:**
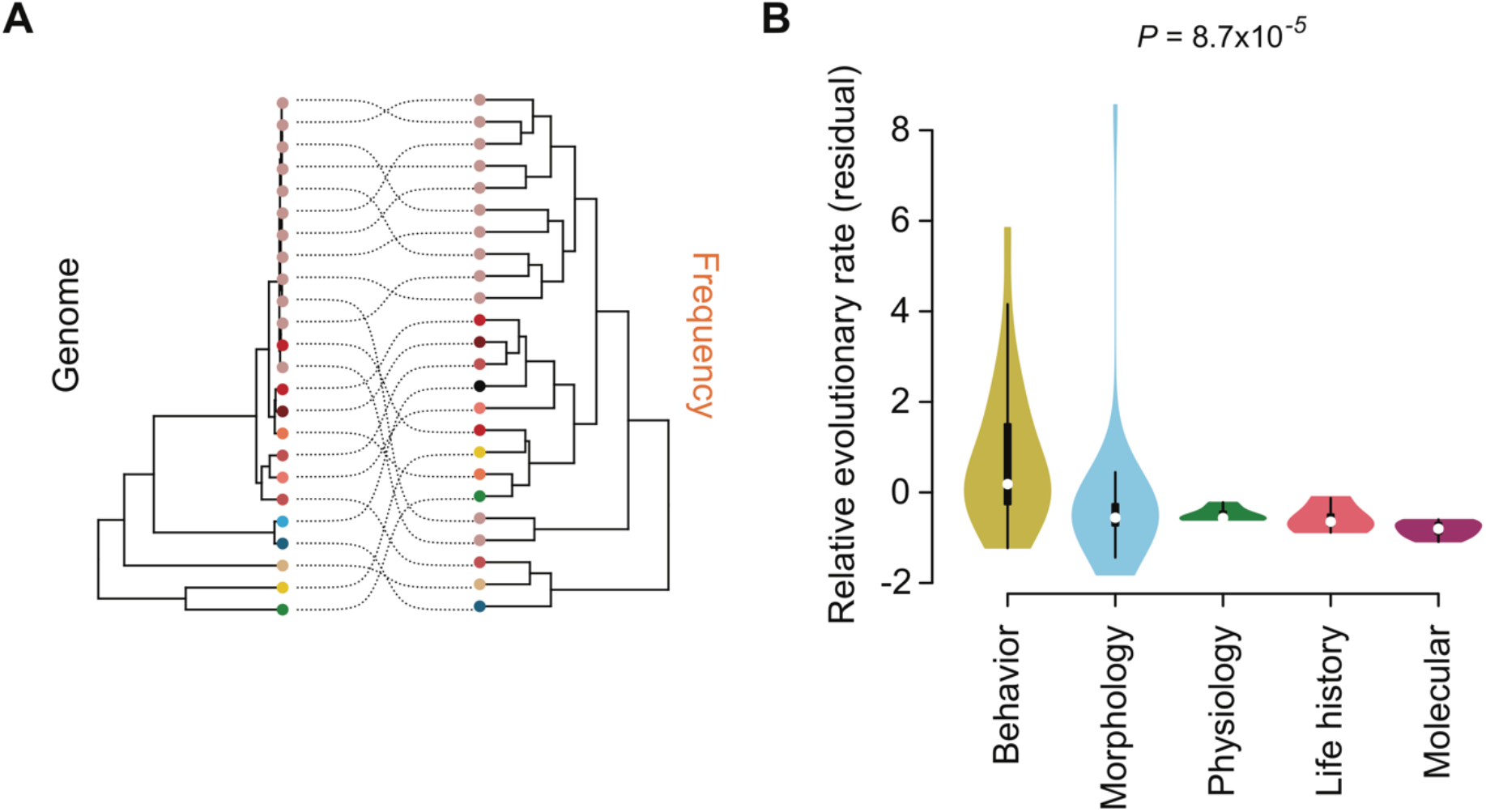
Convergent and rapid evolution of behavioral traits. **(A)** Cophyloplot of *frequency* and the genome phylogeny. Species are represented by colored nodes at the terminal tips of both phylogenies, with their corresponding positions indicated by dotted lines **(B)** Violin plot comparing the residuals of a linear model regressing evolutionary rate on sample size and phylogenetic distance (Kruskal-Wallis test *P*-value).

**Table S1:** Comparing gait Hidden Markov Models. Rows correspond to AIC, BIC, and log likelihood. Columns represent the number of states used to create each HMM modelling the number of legs in stance for all trials tested.

|  | *2 states* | *3 states* | *4 states* | *5 states* |
| --- | --- | --- | --- | --- |
| *AIC* | -7155309 | -7288598 | -7275533 | -7155288 |
| *BIC* | -7155237 | -7288454 | -7275297 | -7154938 |
| *Log Likelihood* | 3577662 | 3644313 | 3637790 | 3577678 |

**Table S2:** Trait meta-analysis data sources. Traits are listed in the left-most column (‘Trait’) followed by the citation (‘Publication’), number of species in the original study that mapped onto the genome phylogeny (‘Number of species’), and the trait category (‘Trait type’).

| *Trait* | *Publication* | *Number of species* | *Trait type* |
| --- | --- | --- | --- |
| *Female locomotion* | Cobb et al. 1986 | 7 | Behavior |
| *Male following* | Cobb et al. 1986 | 7 | Behavior |
| *Wing extension* | Cobb et al. 1986 | 7 | Behavior |
| *Wing rowing* | Cobb et al. 1986 | 7 | Behavior |
| *Scissoring* | Cobb et al. 1986 | 7 | Behavior |
| *Vibrating* | Cobb et al. 1986 | 7 | Behavior |
| *Licking* | Cobb et al. 1986 | 7 | Behavior |
| *Attempted copulation* | Cobb et al. 1986 | 7 | Behavior |
| *Female behavior* | Cobb et al. 1986 | 7 | Behavior |
| *Wing muscle hg1 size male* | Tracy et al. 2020 | 16 | Morphology |
| *Wing muscle hg1 size female* | Tracy et al. 2020 | 16 | Morphology |
| *Wing muscle hg2 size male* | Tracy et al. 2020 | 16 | Morphology |
| *Wing muscle hg2 size female* | Tracy et al. 2020 | 16 | Morphology |
| *Wing muscle hg3 size male* | Tracy et al. 2020 | 16 | Morphology |
| *Wing muscle hg3 size female* | Tracy et al. 2020 | 16 | Morphology |
| *Wing muscle hg4 size male* | Tracy et al. 2020 | 16 | Morphology |
| *Wing muscle hg4 size female* | Tracy et al. 2020 | 16 | Morphology |
| *Wing size female* | Huey et al. 2006 | 10 | Morphology |
| *Wing size male* | Huey et al. 2006 | 10 | Morphology |
| *Thorax size female* | Huey et al. 2006 | 10 | Morphology |
| *Thorax size male* | Huey et al. 2006 | 10 | Morphology |
| *Food preference* | Markow 2015 | 24 | Life history |
| *Dessication resistance female* | Matzkin et al. 2009 | 16 | Physiology |
| *Dessication resistance male* | Matzkin et al. 2009 | 16 | Physiology |
| *Starvation resistance female* | Matzkin et al. 2009 | 11 | Physiology |
| *Starvation resistance male* | Matzkin et al. 2009 | 11 | Physiology |
| *Larval segment 1 position* | Kalay et al. 2020 | 11 | Morphology |
| *Larval segment 2 position* | Kalay et al. 2020 | 11 | Morphology |
| *Larval segment 3 position* | Kalay et al. 2020 | 11 | Morphology |
| *Larval segment 4 position* | Kalay et al. 2020 | 11 | Morphology |
| *Larval segment 5 position* | Kalay et al. 2020 | 11 | Morphology |
| *Larval segment 6 position* | Kalay et al. 2020 | 11 | Morphology |
| *Larval segment 7 position* | Kalay et al. 2020 | 11 | Morphology |
| *Larval segment 8 position* | Kalay et al. 2020 | 11 | Morphology |
| *Female wing display* | Markow & O'Grady 2005 | 12 | Behavior |
| *Number of courtship song* | Markow & O'Grady 2005 | 23 | Behavior |
| *Thermal tolerance female* | Kellerman et al. 2012 | 48 | Physiology |
| *Thermal tolerance male* | Kellerman et al. 2012 | 48 | Physiology |
| *Male thorax length* | Lupold et al. 2016 | 34 | Morphology |
| *Sperm length* | Lupold et al. 2016 | 34 | Morphology |
| *Female thorax length* | Lupold et al. 2016 | 34 | Morphology |
| *Egg volume* | Lupold et al. 2016 | 34 | Morphology |
| *Ovariole number* | Lupold et al. 2016 | 34 | Morphology |
| *Genome size female* | Hjelmen et al. 2019 | 31 | Molecular |
| *Genome size male* | Hjelmen et al. 2019 | 31 | Molecular |
| *Genome size sex difference* | Hjelmen et al. 2019 | 31 | Molecular |
| *Front leg length* | Jezovitz et al. 2020 | 15 | Morphology |
| *Middle leg length* | Jezovitz et al. 2020 | 15 | Morphology |
| *Read leg length* | Jezovitz et al. 2020 | 15 | Morphology |
| *Total leg length* | Jezovitz et al. 2020 | 15 | Morphology |
| *Body size* | Jezovitz et al. 2020 | 15 | Morphology |
| *Ovariole number* | Green & Extavour 2012 | 9 | Morphology |
| *Lifespan female* | Ma et al. 2018 | 13 | Life history |
| *Lifespan male* | Ma et al. 2018 | 13 | Life history |
| *Color spot* | Prudhomme et al. 2006 | 15 | Morphology |
| *Trichome reduction* | Fuqua et al. 2020 | 8 | Morphology |
| *Male pigmentation* | Signor et al. 2016 | 8 | Morphology |
| *Wing display* | Yeh et al. 2006 | 12 | Behavior |
| *Pulse song* | Ding et al. 2019 | 8 | Behavior |
| *Sine song* | Ding et al. 2019 | 8 | Behavior |
| *Mates in dark* | Jezovit et al. 2017 | 27 | Behavior |
| *Environment* | Jezovit et al. 2017 | 46 | Life history |
| *CHC dimorphism* | Jezovit et al. 2017 | 18 | Morphology |
| *Whole-body RNA expression: PC1* | Ma et al. 2018 | 12 | Molecular |
| *Whole-body RNA expression: PC2* | Ma et al. 2018 | 12 | Molecular |
| *Whole-body RNA expression: PC3* | Ma et al. 2018 | 12 | Molecular |
| *Transitions: PC1* | This study | 24 | Behavior |
| *Transitions: PC2* | This study | 24 | Behavior |
| *Frequency: PC1* | This study | 24 | Behavior |
| *Frequency: PC2* | This study | 24 | Behavior |
| *Structure* | This study | 24 | Behavior |
| *Louvain cluster 1 frequency* | This study | 24 | Behavior |
| *Louvain cluster 2 frequency* | This study | 24 | Behavior |
| *Louvain cluster 3 frequency* | This study | 24 | Behavior |
| *Louvain cluster 4 frequency* | This study | 24 | Behavior |
| *Louvain cluster 5 frequency* | This study | 24 | Behavior |
| *Louvain cluster 6 frequency* | This study | 24 | Behavior |
| *Louvain cluster 7 frequency* | This study | 24 | Behavior |

